# The importance of Indigenous Peoples’ lands for the conservation of terrestrial vertebrates

**DOI:** 10.1101/2019.12.11.873695

**Authors:** Christopher J. O’Bryan, Stephen T. Garnett, John E. Fa, Ian Leiper, Jose Rehbein, Álvaro Fernández-Llamazares, Micha V. Jackson, Harry D. Jonas, Eduardo S. Brondizio, Neil D. Burgess, Catherine J. Robinson, Kerstin K. Zander, Oscar Venter, James E.M. Watson

## Abstract

Indigenous Peoples’ lands cover over one-quarter of the Earth’s surface, a significant proportion of which is still free from industrial-level human impacts. As a result, Indigenous Peoples’ lands are crucial for the long-term persistence of Earth’s biodiversity and ecosystem services. Yet, information on species composition within Indigenous Peoples’ lands globally remains unknown. Here, we provide the first comprehensive analysis of terrestrial vertebrate composition across mapped Indigenous lands by using distribution range data for 20,328 IUCN-assessed mammal, bird and amphibian species. We estimate that 12,521 species (62%) have ≥10% of their ranges in Indigenous Peoples’ lands, and 3,314 species (16%) have >half of their ranges within these lands. For threatened species assessed, 1,878 (41.5% of all threatened of all threatened mammals, birds and amphibians) occur in Indigenous Peoples’ lands. We also find that 3,989 species (of which 418 are threatened) have ≥10% of their range in Indigenous Peoples’ lands that have low human pressure. Our results are conservative because not all known Indigenous lands are mapped, and this analysis shows how important Indigenous Peoples’ lands are for the successful implementation of international conservation and sustainable development agendas.

## 1. INTRODUCTION

Through well-established traditional knowledge systems and governance practices, Indigenous Peoples are the environmental stewards of their lands. This is gradually being recognized in domestic and international policy (Brondizio et al. 2019). A recent analysis indicates that Indigenous Peoples’ lands cover at least a quarter of terrestrial Earth, overlapping with 37% of all terrestrial protected areas and with 40% of landscapes without industrial-level human impacts (Garnett et al. 2018). Some countrywide assessments demonstrate the importance of Indigenous Peoples’ lands in terms of the biodiversity contained within them. In Australia, for example, 45-60% of the country’s threatened species are found in Indigenous Peoples’ lands (Renwick et al. 2017; Leiper et al. 2018) and vertebrate biodiversity in Indigenous People’s lands in three countries (Australia, Brazil and Canada) are comparable to those found in protected areas (Schuster et al. 2019). However, global assessments of the overlap between Indigenous Peoples’ lands and species distributions (including threatened species) are lacking, especially the relationship with areas free from industrial-level human impacts. Such regions are likely to be of high conservation value (Di Marco et al. 2018), given the connection between land use transformation and species declines (Newbold et al. 2015; Maxwell et al. 2016; Tilman et al. 2017). These landscapes may also be important ecological refugia (Scheffers et al. 2016; Allan et al. 2019) offering some protection against the pressures of expanding resource extraction frontiers.

Here, we provide the first global assessment of the overlap between mapped Indigenous Peoples’ lands (Garnett et al. 2018) and terrestrial vertebrate ranges (IUCN 2016; Birdlife International and Handbook of the Birds of the World 2017). We also assess species composition within Indigenous Peoples’ lands with low human pressure using updated ‘Human Footprint’ data (Venter et al. 2016). These results are relevant to the development and implementation of the post-2020 Global Biodiversity Framework agreement that will emerge from Convention for Biological Diversity’s (CBD) target discussions on abating species extinctions and reducing the erosion of ecosystem services (Watson & Venter 2017; CBD 2018) as well as for nations trying to implement actions to achieve the 2030 United Nation’s Sustainable Development Goals.

## 2. METHODS

### Species distribution data

We focused our analysis on the terrestrial vertebrates (mammals, amphibians and birds) that had been comprehensively assessed by the IUCN by 2017. Spatial data on mammal and amphibian distributions were obtained from the IUCN Red List of Threatened Species (IUCN 2017), and bird distributions from BirdLife International (Birdlife International and Handbook of the Birds of the World 2017). We excluded species considered Extinct, Data Deficient, or any other extant native and reintroduced species whose distribution did not intersect with the spatial datasets employed in this study.

### Spatial data on Indigenous lands

Globally, more than 370 million persons in more than 70 countries self-identify as Indigenous Peoples (ILO 2019; see Garnett et al. 2018 for discussion of definitions). We used recently-compiled global spatial data on Indigenous Peoples’ lands, sourced or delineated on the basis of open-access published sources (see Garnett et al. 2018 for further details) that, while certainly incomplete, is the best available spatial layer.

### Spatial data on human pressure

Advances in remote sensing coupled with bottom-up survey data, have enabled the development of a spatially explicit, validated, high-resolution global dataset on human pressures (Venter et al. 2016a). These datasets permit the quantification of the extent of intense pressures on individual species (Di Marco et al. 2018; Allan et al. 2019). We used the most current Human Footprint map available (from 2013), comprising a composite spatial index of key human pressures on natural ecosystems at a 1-km^2^ resolution.

Eight human pressure variables were used in the Human Footprint: 1) built environments, 2) population density, 3) electric infrastructure, 4) crop lands, 5) pasture lands, 6) roads, 7) railways, and 8) navigable waterways. These eight individual pressures are scaled between 0 and 10 based on their estimated environmental impact and summed in 1km^2^ grid cells. Some pressures co-occur whilst others are mutually exclusive, resulting in a combined global scale between 0 and 50 where 0 has no detectable change and 50 is extreme urban conglomerates. We reclassified the Human Footprint map to an index threshold of < 3 because this threshold is now considered the standard for evaluating the degree of low human pressure across ecosystems (Di Marco et al. 2018; Jones et al. 2018). A threshold of ∼3 is where areas with low states of human pressure transition to human-dominated activities such as pastureland (Venter et al. 2016b). Importantly, index values at or >3 reveal an increased extinction risk in mammals (Di Marco et al. 2018).

### Analysis

We combined the spatial datasets on Indigenous Peoples’ lands and low-Human Footprint data (not including species data) in a single spatial data layer (in raster format) using a World Molleweide projection in a geographic information system (ESRI ArcGIS) at a 1 km^2^ resolution. To do this, we converted each feature polygon layer into raster format and reclassified each layer with a layer-explicit value. We summed each raster layer using the raster calculator tool, which resulted in a single raster image containing the values for each layer within each cell. For example, the cells within this single raster image contained information on the presence of Indigenous Peoples’ lands (Garnett et al. 2018) and whether the cell contained landscapes with low human footprint (Venter et al. 2016a, 2016b). The layer was then intersected with the ranges of 20,328 terrestrial vertebrates - 4,592 mammal species, 10,743 bird species, and 4,993 amphibian species. We calculated the proportion of land occupied by each species in Indigenous Peoples’ lands, in low-pressure landscapes or in both using the tabulate area tool within ArcGIS 10.5 Model Builder (ESRI 2017) and R statistical software (R Core Team 2017). The proportions of all species and threatened species on each land use type were compared using Chi Square with Yates’ correction (Yates 1934).

## 3. RESULTS

### Occurrence of species in Indigenous Peoples’ lands

Indigenous Peoples’ lands encompass a total of at least 38 million km^2^ (28.3 %) of terrestrial Earth (Garnett et al. 2018; Table S1). We find that 12,521 (61.6%) of all species assessed have at least 10% of their ranges within these lands, and 3,314 (16.3%) have >50% of their range in these lands (Table 1). Birds have a higher percentage of species with at least 10% of their ranges in Indigenous People’s lands than other classes: 69.6% (n = 7,481) compared to 60.8% (n = 2,791) of mammals and 45.0% (n = 2,249) of amphibians. Birds also have the highest mean range proportion containing these lands (25.8% of range on average) followed by mammals (25.0%) and amphibians (21.8%) (Figure 1). However, amphibians have a greater percentage of species with >50% of their ranges in these lands: 17.8% (n = 889) of amphibians versus 17.6% (n = 807) of mammals and 15.1% (n = 1,618) of birds (Table 1). Australia and Indonesia have the highest proportion of species with >50% of their range within Indigenous Peoples’ lands with east Asia, Africa and the mountainous central and South America also having high proportions (Figure 2).

**Table 1.**
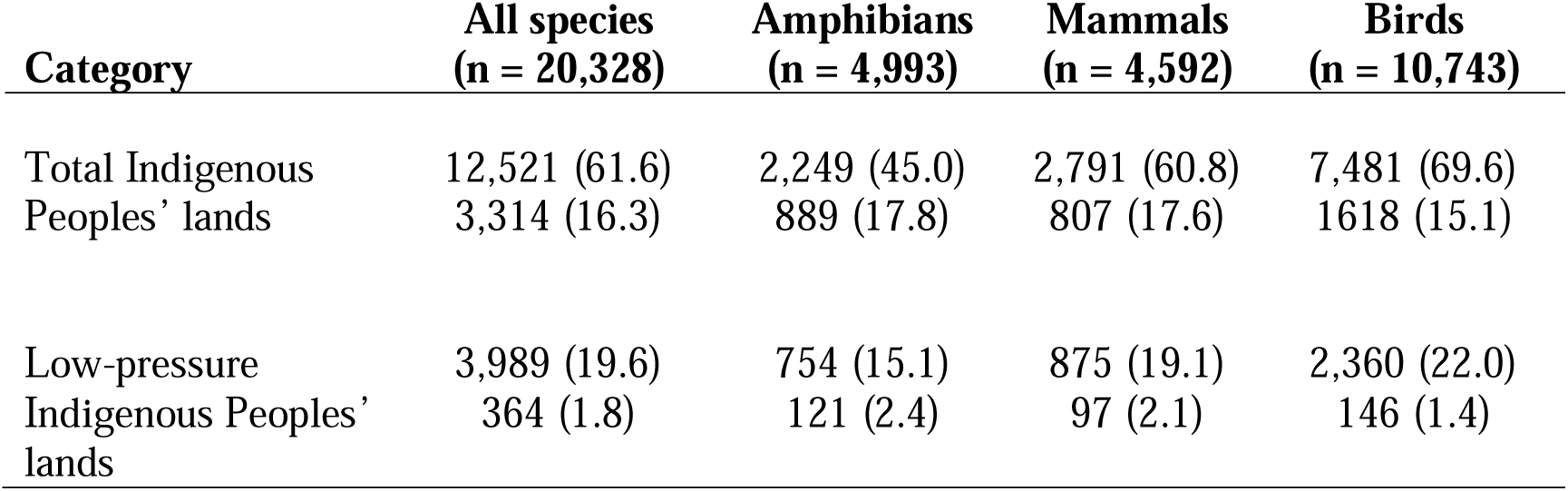
The number and percentage of species that have ≥10% (top number) and >50% (bottom number) of their range overlapping with a specified land category.

| **Category** | **All species**<br>**(n = 20,328)** | **Amphibians**<br>**(n = 4,993)** | **Mammals**<br>**(n = 4,592)** | **Birds**<br>**(n = 10,743)** |
| --- | --- | --- | --- | --- |
| Total Indigenous Peoples' lands | 12,521 (61.6)<br>3,314 (16.3) | 2,249 (45.0)<br>889 (17.8) | 2,791 (60.8)<br>807 (17.6) | 7,481 (69.6)<br>1,618 (15.1) |
| Low-pressure Indigenous Peoples' lands | 3,989 (19.6)<br>364 (1.8) | 754 (15.1)<br>121 (2.4) | 875 (19.1)<br>97 (2.1) | 2,360 (22.0)<br>146 (1.4) |

**Figure 1.**
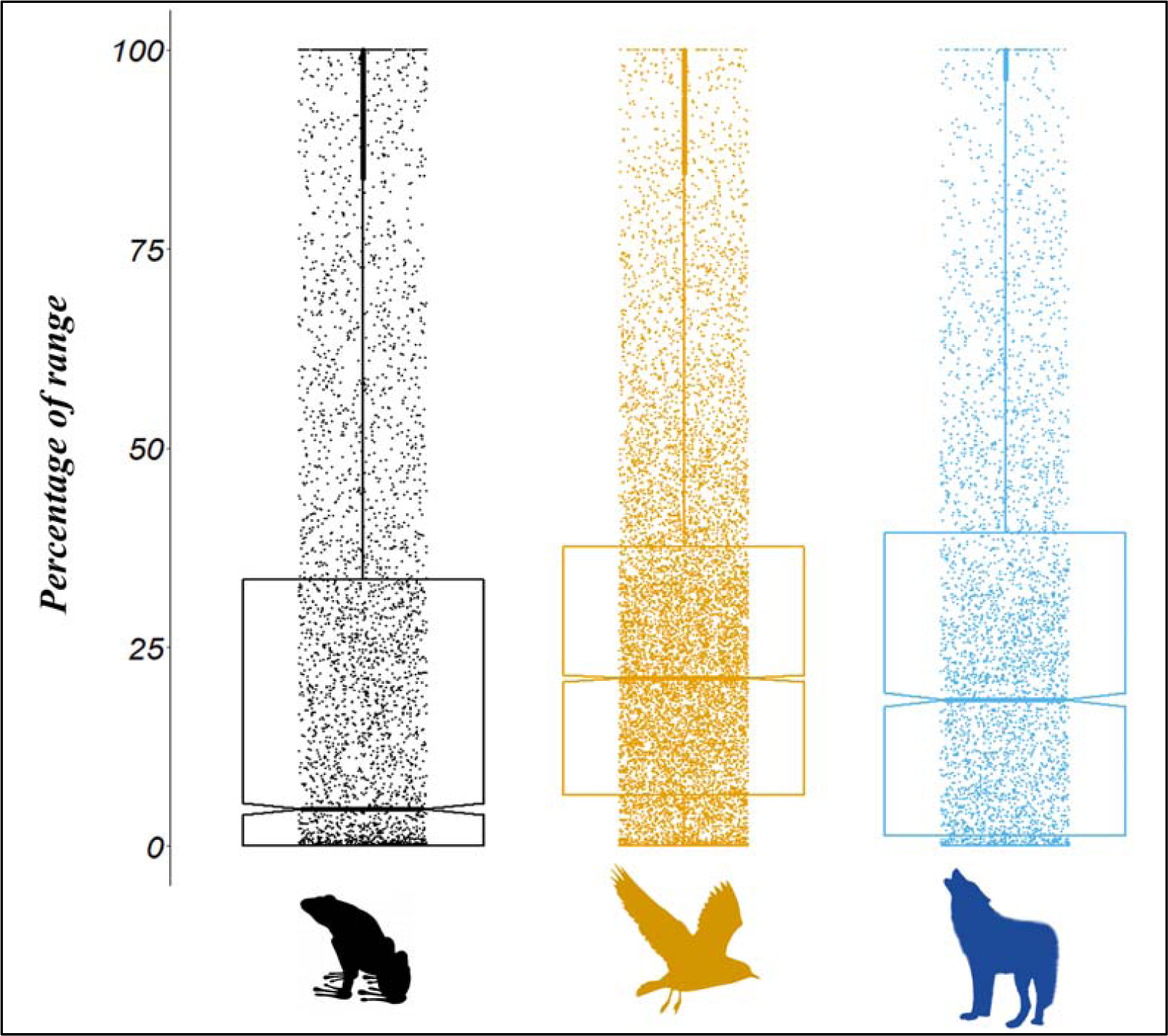
The percentage of species’ ranges containing Indigenous Peoples’ lands across amphibians (n = 4,993), birds (n = 10,743), and mammals (n = 4,592) assessed by the IUCN Red List. Each dot represents an individual species. The box plot denotes the lower quartile, mean, and upper quartile for each species class.

**Figure 2.**
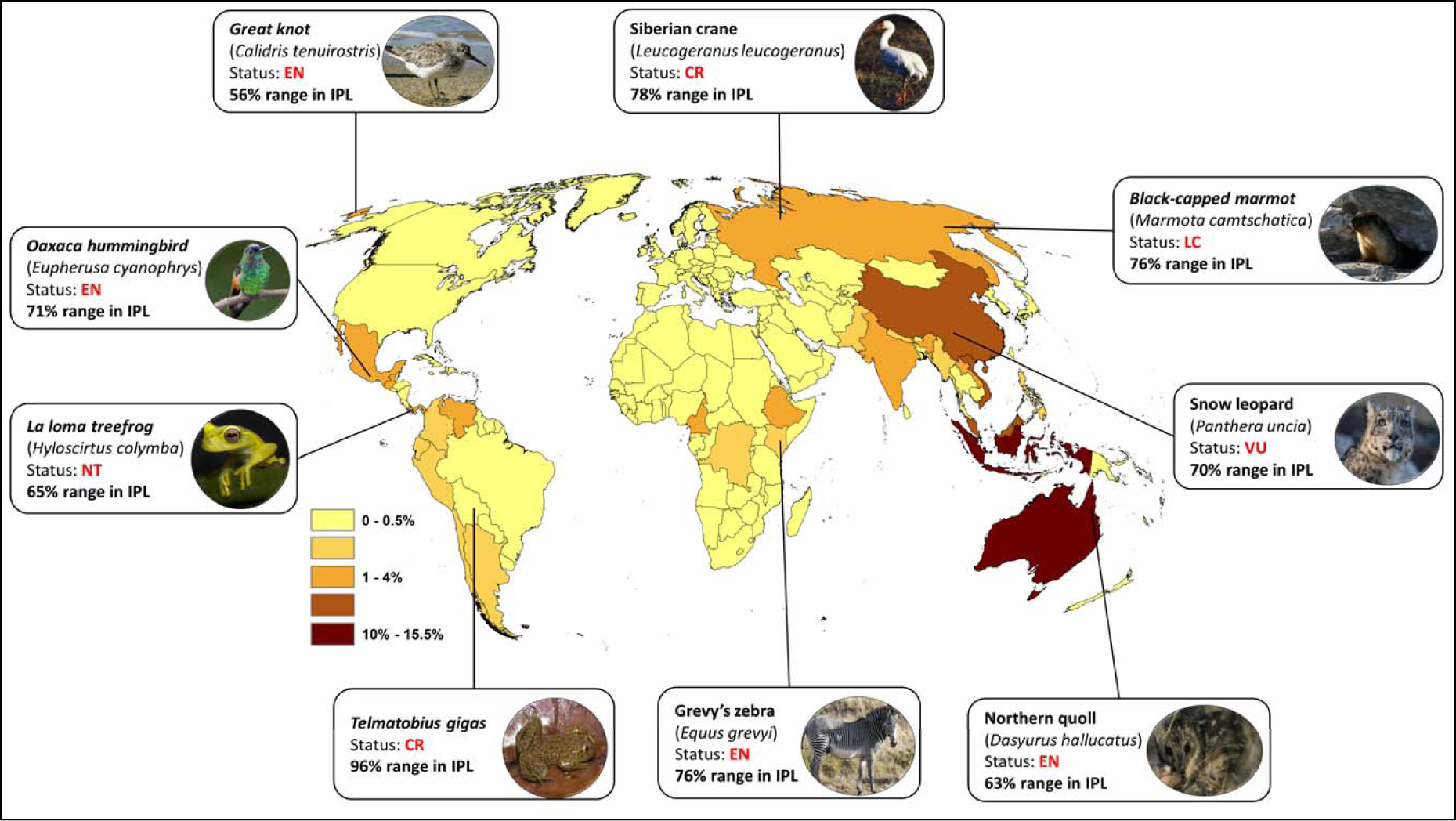
The proportion of species within each country that have >50% of their range in Indigenous Peoples’ lands (IPL) that are mapped Garnett et al. 2018), with a subset of exemplar species.

Of the 4,523 species classified as threatened (i.e. at least Vulnerable on the IUCN Red List), 1,878 (41.5%) have at least 10% of their ranges within Indigenous Peoples’ lands, with 952 (50.7%) being Vulnerable, 637 (33.9%) being Endangered, and 289 (15.4%) being Critically Endangered on the IUCN Red List. We also find that 840 (18.6%) of all threatened species have >50% of their ranges within these lands (Table 2), with 393 (46.8%) being Vulnerable, 268 (31.9%) Endangered, and 179 (21.3%) Critically Endangered species. The proportion of threatened species with >50% of their range on Indigenous Peoples’ lands was higher than for all species (χ^2^ with Yates’ correction 6.86, P<0.01).

**Table 2.**
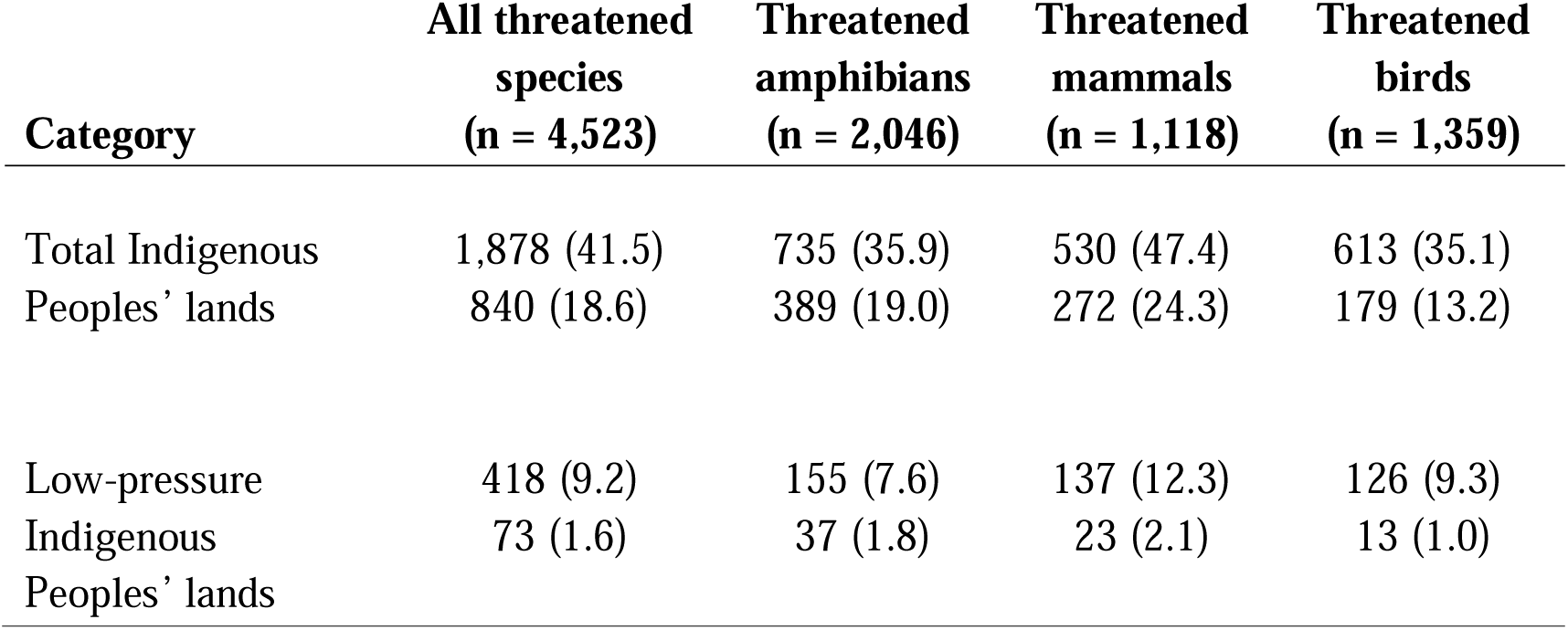
The number and percentage (in parentheses) of threatened species that have ≥10% (top number) and >50% (bottom number) of their range overlapping with a specified land category.

We find that threatened amphibians have a slightly higher percentage of species with at least 10% of their ranges in Indigenous Peoples’ lands than other threatened classes: 35.9% (n = 735) compared to 35.1% (n = 613) of birds and 47.4% (n = 530) of mammals. However, threatened mammals have a greater percentage of species with >50% of their range in these lands: 24.3% (n = 272) versus 19.0% (n = 389) of threatened amphibians and 13.2% (n = 179) of threatened birds (Table 2).

### Occurance of species in Indigenous Peoples’ lands with low human pressure

Nearly 21 million km^2^ of Indigenous Peoples’ lands (15.5% of all terrestrial Earth, and 45.2% of all Indigenous Peoples’ lands) have low human pressure (Table S1). We find that 3,989 (19.6%) of species assessed have at least 10% of their range in these low-pressure Indigenous Peoples’ lands, with 364 (1.8%) having >50% of their ranges in these lands (Table 1). Birds have a higher percentage of species with at least 10% of their ranges in Indigenous Peoples’ lands with low human pressure than other classes: 22.0% (n = 2,360) compared to 19.1% (n = 875) of mammals and 15.1% (n = 754) of amphibians. Still, amphibians have a greater percentage of species with >50% of their ranges in Indigenous Peoples’ lands with low human pressure: 2.4% (n = 121) versus 2.1% (n = 97) of mammals and 1.4% (n = 146) of birds.

We find that the percentage of threatened species within low-pressure Indigenous Peoples’ lands is considerably lower than that of threatened species across all Indigenous Peoples’ lands (Figure 3). As many as 418 (9.2%) of the threatened species assessed have at least 10% of their ranges in low-pressure Indigenous Peoples’ lands, with 240 (57.4%) of these being Vulnerable, 117 (28.0%) Endangered, and 61 (14.6%) Critically Endangered species. We also estimate that 73 (1.6%) of the threatened species assessed have >50% of their ranges in these lands (Table 2), with 48 (65.8%) of these being Vulnerable, 14 (19.2%) Critically Endangered, and 11 (15.1%) Endangered species (Figure 3B). The proportion of threatened species with >50% of their range on low-pressure Indigenous Peoples’ lands did not differ significantly from the proportion for all species (χ^2^ with Yates’ correction 0.798, P>0.10).

**Figure 3.**
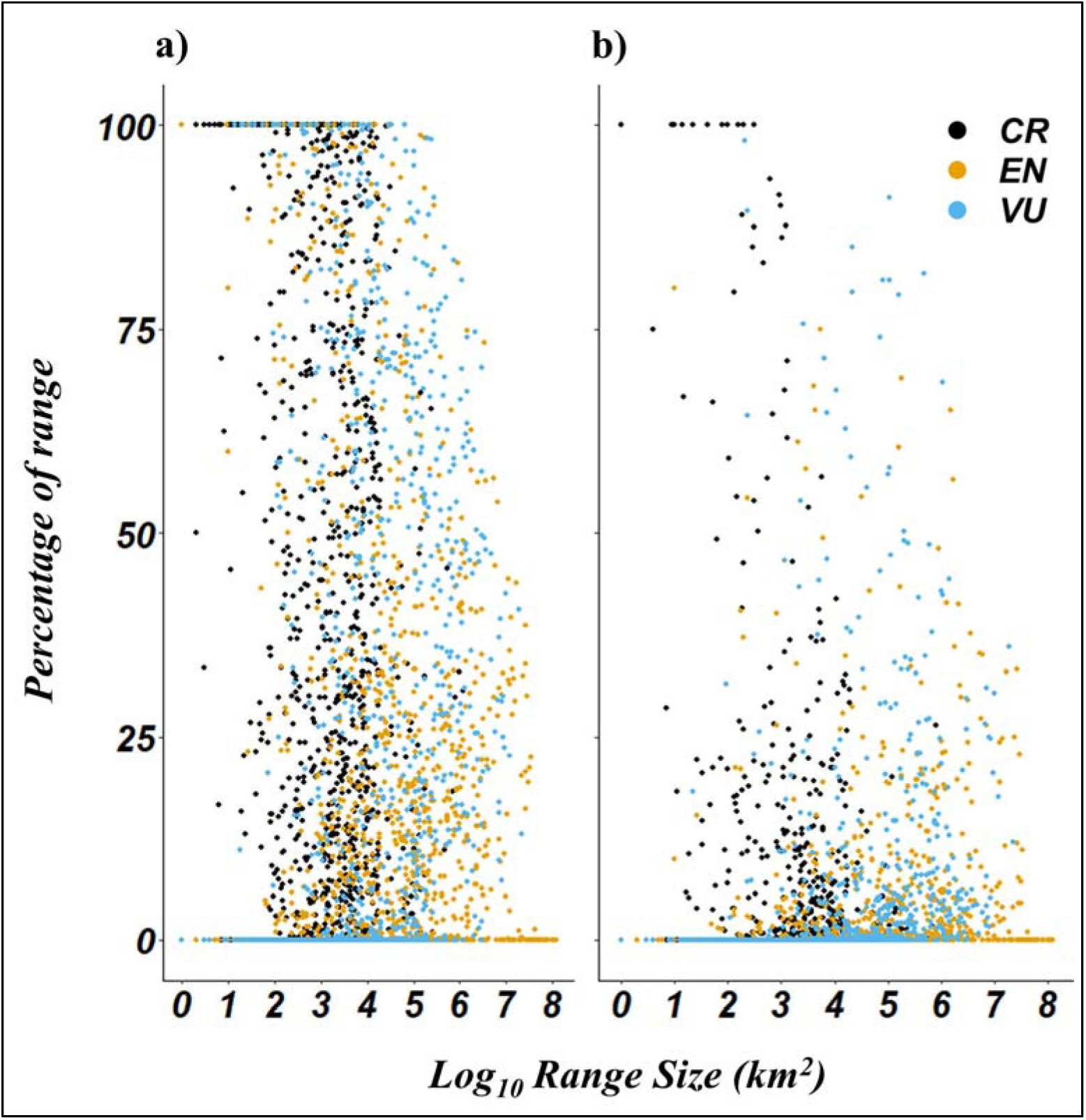
The proportion of threatened species ranges (a) within Indigenous Peoples’ lands and (b) Indigenous Peoples’ lands that are mapped (Garnett et al. 2018) with low human pressure broken down by levels of extinction risk as designated by the IUCN (CR = Critically Endangered; EN = Endangered; VU = Vulnerable).

We estimate that threatened mammals have a higher percentage of species with at least 10% of their ranges in low-pressure Indigenous Peoples’ lands than other threatened classes: 12.3% (n = 137) compared to 9.3% (n = 126) of birds and 7.6% (n = 155) of amphibians (Table 2). Similarly, threatened mammals have a greater percentage of species with >50% of their range in these lands: 2.1% (n = 23) versus 1.8% (n = 37) of threatened amphibians and 1.0% (n = 13) of threatened birds (Table 2).

## 4. DISCUSSION

Indigenous Peoples’ lands cover a large portion of Earth’s land surface (Garnett et al. 2018), and also include some of the highest quality forest lands worldwide (Fa et al. in press). It follows that Indigenous Peoples are also custodians of a substantial proportion of the world’s biodiversity. While it has long been suspected that the proportion of biodiversity in Indigenous Peoples’ lands was likely to be high (Toledo 2013), our study is the first to use robust, repeatable methods for determining this. The numbers we have derived are substantial: globally 62% of mammals, birds and amphibians have ≥10% of their range within Indigenous Peoples’ lands; for 16% of species, including 18% of threatened species, the proportion is >50%.

These figures are conservative for a number of reasons. The taxonomic range of the species groups for which we have range data – mammals, birds and amphibians – is but a small fraction of the biodiversity found (Larsen et al. 2017). Our results, based on best available globally consistent species data, is also likely to be true of plants, invertebrates and others forms of biodiversity that could increase the absolute number of species by several orders of magnitude (but see Oberprieler et al. 2019). Moreover, because stringent legislation often controls access to and activities within Indigenous Peoples’ lands, affecting the extent to which biodiversity is documented and mapped (dos Santos et al. 2015), it is very likely that survey effort in these lands is still incomplete (e.g., Bernard et al. 2011 for bats in the Amazon). A corollary of this is that Indigenous Peoples could potentially play an important role in filling species knowledge gaps and information deficits in large parts of the planet, as well as in furthering our understanding of the conservation status and population trends of many species, but also levels of species richness, endemism hotspots and largely unknown distribution range discontinuities (Johnson et al. 2015). Indeed, 42.4% (935 out of 2,203) of Data Deficient species from our dataset have ≥10% of their ranges within Indigenous Peoples’ lands. We also note that many of areas not yet mapped as Indigenous Peoples’ lands are also likely to retain an Indigenous connection so our numbers are an under-estimate of overall species coverage.

Myriad examples are available on how collaboration between Indigenous Peoples and researchers has refined knowledge of species ecological distribution ranges, baselines and trends (e.g. Mistry & Berardi 2016; Skroblin et al. 2019). However, such knowledge partnerships need to be negotiated appropriately (Robinson et al. 2016). The central message from this analysis, that Indigenous Peoples’ participation, lands and perspectives are vital to any policies and programs aiming to further global biodiversity conservation, strongly aligns with that of the Intergovernmental Science-Policy Platform on Biodiversity and Ecosystem Services (Brondizio et al. 2019; Diaz et al. 2019; IPBES 2019) and many other studies (e.g. Dinerstein et al. 2019; Reyes-García et al. 2019).

Our results point to the fact that Indigenous Peoples’ rights must be fully respected, including their full and effective participation in developing laws, policies and programs that affect them. Although representatives of Indigenous Peoples are gradually engaging in global environmental governance, such as IPBES, the Intergovernmental Panel on Climate Change and the CBD, this often occurs in the face of substantial barriers to engagement related to scale, knowledge and power (Brugnach et al. 2017). Greater recognition and support for the relationships that Indigenous Peoples have with their lands and their natural resources is therefore a pressing imperative from the perspective of both social equity and biodiversity conservation (Howitt 2018). Only through equitable partnerships and other forms of collaboration with fully empowered Indigenous Peoples will it be possible to discuss, but more importantly ensure the long-term conservation of biodiversity on Indigenous Peoples’ lands.

## Acknowledgments

The work was partially funded by the NASA Biodiversity and Ecological Forecasting Program under the 2016 ECO4CAST solicitation through grant NNX17AG51G. JEF was funded by the US Agency for International Development as part of the Bushmeat Research Initiative of the CGIAR research program on Forests, Trees and Agroforestry.

