## Supplementary Material for "The importance of Indigenous Peoples’ lands for the conservation of terrestrial vertebrates"

**This file contains: 1 Table**

| **Land Categories** | **Area (km^2^)** | **Proportion of Earth’s terrestrial surface** | **Source** |
| --- | --- | --- | --- |
| Total terrestrial surface | 134,154,306 | 1.0000 |  |
| Total Indigenous Peoples’ lands | 38,001,845 | 0.2833 |  |
| Pressure-free Indigenous Peoples’ lands | 20,825,442 | 0.1552 |  |
| Formally protected Indigenous Peoples Lands | 7,077,09 | 0.02666 |  |

**Table S1.** The area of each land category used for our study, and the proportion of each on Earth’s terrestrial surface.

Data availability

Species distribution data are available for download from the IUCN Spatial Data Download centre (IUCN 2017). Data used for Indigenous Peoples’ land mapping are provided in Supplementary Information section of Garnett and colleagues (Garnett et al. 2018) and the derived maps are available from the author S.T.G. on reasonable request. The protected areas data that support the findings of this study are available from the UN Environment World Conservation Monitoring Centre (UNEP-WCMC & IUCN 2016), the Human Footprint data are available in the Dryad Digital Repository (Venter et al. 2016).
